# Rapid emergence of clonal interference during malaria parasite cultivation

**DOI:** 10.1101/2020.03.04.977165

**Authors:** Catherine Jett, Aliou Dia, Ian H. Cheeseman

## Abstract

Laboratory cultivation of the malaria parasite *Plasmodium falciparum* has underpinned nearly all advances in malariology in the past 30 years. When freshly isolated clinical isolates are adapted to *in vitro* culture mutations rapidly fix increasing the parasite growth rate and stability. While the dynamics of culture adaptation are increasingly well characterized, we know little about the extent of genomic variation that arises and spreads during long term culture. To address this we cloned the 3D7 reference strain and maintained a culture for ~84 asexual cycles (167 days). Growth rate of the culture population increased 1.14-fold over this timeframe. We used single cell genome sequencing of parasites at cycles 21 and 84 to measure the accumulation of diversity *in vitro*. This parasite population showed strong signals of adaptation across this time frame. By cycle 84 two dominant clades had arisen and were segregating with the dynamics of clonal interference. This highlights the continual process of adaptation in malaria parasites, even in parasites which have been extensively adapted to long term culture.

## Introduction

The ability to maintain long-term cultures of *Plasmodium falciparum* has been one of the most fundamental advances in malaria research (Trager and Jensen, 2005, Trager and Jensen, 1997, Trager and Jensen, 1976). Parasite culture is so pervasive it is challenging to find advances that have not been enabled by it. During *in vitro* culture malaria parasites are maintained in a suspension of red blood cells (RBCs) obtained from a donor, supplied with the necessary nutrients from culture media and serum and housed in a low oxygen environment at normal body temperature (37C°). By regular changes in media and replenishing RBCs parasite cultures can be maintained indefinitely, and subject to drug testing, genetic manipulation (Ghorbal et al., 2014), and placed under selective pressures to understand adaptation (Cowell et al., 2018).

Parasites isolated from malaria infected patients often replicate poorly during early passages *ex vivo*. While the precise mechanisms underlying this are elusive, a number of investigators have catalogued genetic changes occurring during the early phase of culture adaptation. These include the loss or gain of large chromosomal regions (Biggs et al., 1989, Carret et al., 2005, Nair et al., 2010), and specific site mutations (Claessens et al., 2017). A common theme tying many of these alterations together is the process of gametocytogenesis. In malaria-infected patients, the malaria parasite must commit a small proportion of parasites to sexual stage development. Mature gametocytes are then taken up during a mosquito blood meal and transmitted to future hosts. However, *in vitro* the parasite only requires cycling through the asexual stages. The lack of pressure to maintain commitment to sexual stages (or some biochemical benefit in shutting off these pathways) is thought to be the underlying mechanism.

Long-term adaptation experiments in yeast and bacteria have revealed that adaptation to a novel environment is not limited (Barrick and Lenski, 2013, Barrick et al., 2009, Good et al., 2017, Lang et al., 2013). Over the course of its 32 year, ~70,000 generation duration the long term evolution experiment has revealed that novel adaptive mutations are continually arising in bacteria, albeit with diminishing returns on increased fitness (Arjan et al., 1999). Experimental evolution over such long time frames has shown the dynamics of adaptation can be complex. Beneficial mutations may arise either on different genetic backgrounds within a culture, or sequentially on the same genetic background. The former of these leads to competition between lineages bearing different adaptive mutations as both rise in frequency (clonal interference) and the latter to waves of adaptation where single mutations are swept through a population in succession (Barrick and Lenski, 2013).

Mutation accumulation experiments have directly measured the mutation rate of the parasite genome (~3×10^−10^ genome^−1^ lifecycle^−1^ for SNPs (Bopp et al., 2013, Claessens et al., 2014, Hamilton et al., 2017, McDew-White et al., 2019)). This means in a 10ml malaria parasite culture (1% parasitaemia, 2% haematocrit) there will be >70,000 point mutations every 48 hours. The supply of mutations is only one element of adaptation. Each of novel mutations is initially at an extraordinary low frequency (~1 in 1×10^−7^ parasites). At this frequency even strongly beneficial mutations are lost by genetic drift most of the time. This dynamic has been exploited to understand the evolution of antimalarial drug resistance *in vitro* (Cowell et al., 2018). It becomes much easier for drug resistance mutations to arise when all sensitive parasites are eliminated by drug treatment.

We expect the diversity present in a culture to be shaped by the processes of mutation, selection and drift. However, we know little about how often beneficial mutations emerge and spread in even extraordinary well studied systems such as culture adapted parasite lines. To address this we performed single cell sequencing of a long term cultured parasite line and measured clonal dynamics over time.

## Results

### Long term cultivation of 3D7 increases growth rate

We cloned the 3D7 reference strain by limiting dilution, and allowed the culture to expand to a 5% parasitaemia over 21 days, we refer to this as day 0. Every ~48 hours we measured parasitaemia and sub-cultured parasites to 0.5−1.0% parasitaemia. By measuring the increase in parasitaemia over the subsequent 2 days (Figure 1A) we estimated the growth rate of the population (Figure 1B). Over 167 days (83.5 life cycles) we saw an increase in growth rate by 1.14 fold (*p*=0.025, linear model). The culture was stable and healthy over the duration of the experiment despite peaks in parasitaemia of over 12%.

### Characterizing diversity by single cell sequencing

**Figure 1.**
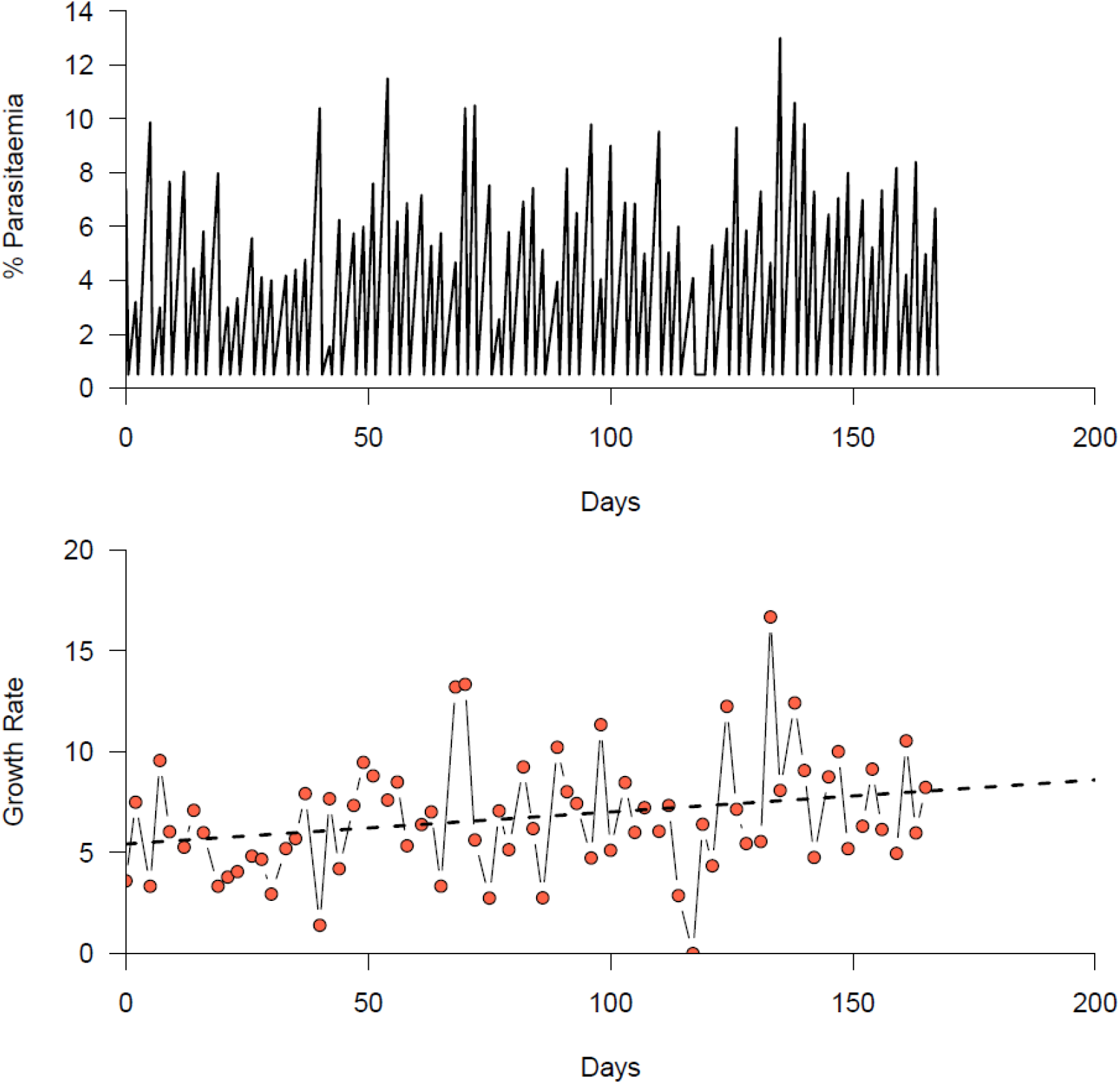
Parasite growth over 140 days. (A) Parasitaemia measured from giemsa stained slides over the duration of the experiment. After each measurement cultures were reduced to 0.5% parasitaemia. (B) Growth rate per lifecycle. Dashed line represents a linear model fit to the data to estimate the change in growth rate over time.

We performed single cell sequencing of 20 cells each from day 56 and day 140 using the method of Trevino et al (Trevino et al., 2017) and bulk sequencing of the population at days 0, 42, 84 and 140. This generated high quality data for 32 cells, with at least 5 reads mapping to 60% of the genome. After filtering the data to exclude mutations present in the day 0 time point, we detected 24 high quality mutations in the data set, including 12 SNPs and 12 indels (Table 1). We did not include amplifications in our survey of mutations as these are not called with high confidence in our data. At day 56 we saw an average of 0.47 mutations per line (0.11 SNPs, 0.37 indels). At day 140 we saw an average of 2.65 mutations per line (1.35 SNPs, 1.3 indels). Published mutation rate suggest a parasite at day 56 will have accrued an average of 0.2 SNPs, 1.36 indels, and a parasite at day 140 will have accrued an average of 0.5 SNPs, 3.40 indels in the absence of selection.

### Phylogenetic reconstruction of single cell sequences reveals clonal dynamics

Using the SNP data we reconstructed the phylogenetic relationship between individual cells using BEAST (Figure 2). We inferred a time stamped tree including information on the sampling date for each cell.

Bayesian Skyline plots showed the effective population size of the culture increased linearly over time as more mutations were introduced into the culture. There was a plateau in the increasing population size after day 56 to day 90. This may be the result of adaptive mutations sweeping through the population.

**Figure 2.**
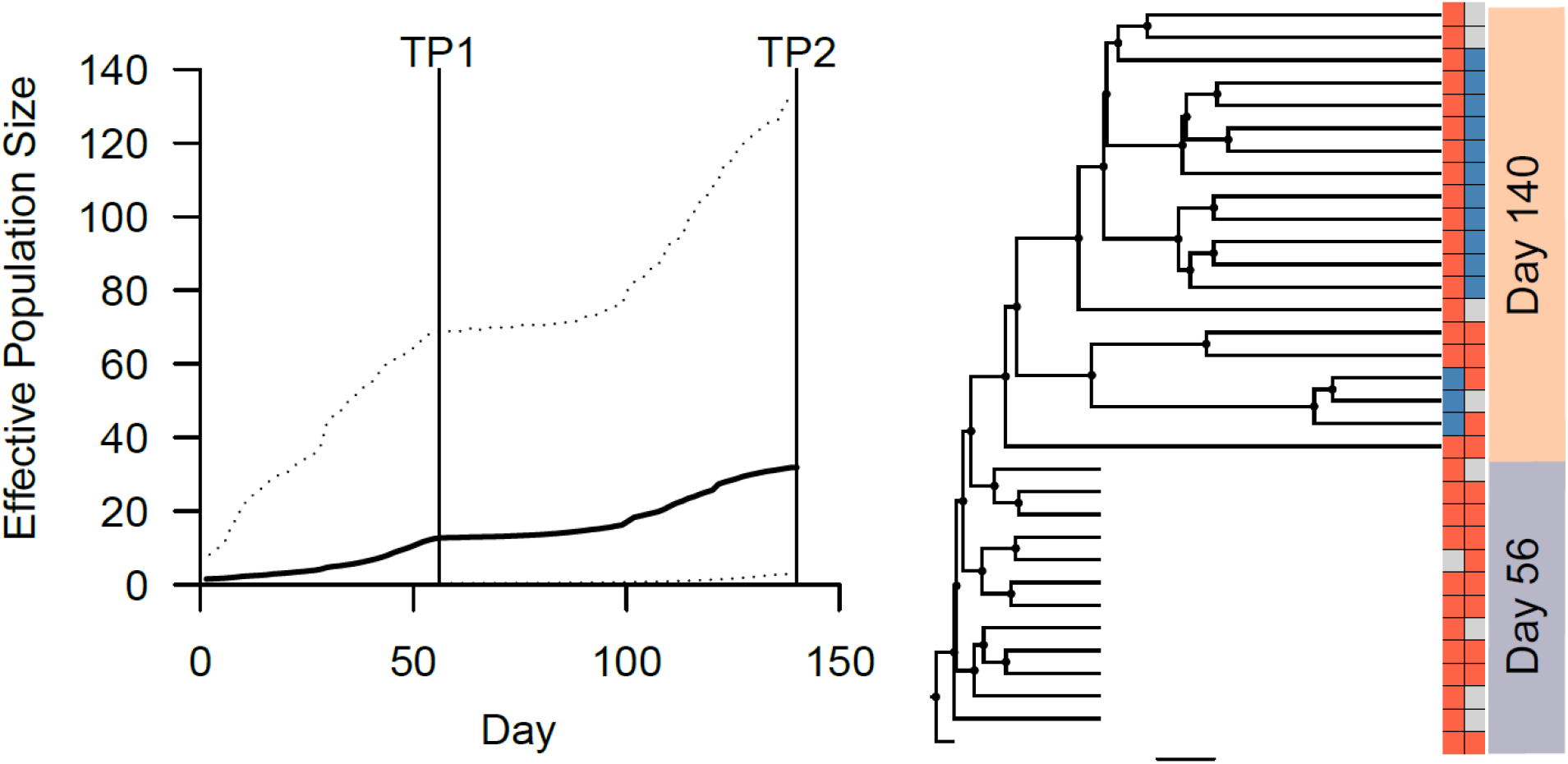
Phylogenetic reconstruction of a malaria parasite culture. (A) Bayesian Skyline plot of effective population size over time. The two sampling timepoints of the single cell sequence data are shown by vertical black lines labeled TP1 and TP2. Dashed lines show the 95% confidence intervals. (B) A time-stamped tree reconstructed from SNP data. To the right of the tree a matrix showing the driver mutations is shown. Blue boxes denote the derived allele and red the ancestral allele, missing data is shown by a grey box. The sampling time for each cell is shown on the right of this matrix.

### Adaptive mutations

We find two SNP mutations where their increase in frequency indicates they are highly adaptive. The first introduces a premature stop codon (AAA > TAA) at amino acid position 693 of PF3D7_1325800, a conserved protein. The second is a non-synonymous mutation at position 1151 changing an alanine to a glutamic acid (GCA > GAA) in PF3D7_1146600, oocyst-rupture protein 1 (ORP1). We estimated their frequency from bulk genome sequencing across the duration of the experiment where they rose from 0% to 31.1% and 53.9% respectively. Based on their trajectory we estimate selection coefficients to be 0.17 and 0.16.

Positive selection may be acting upon many mutations detected in our analysis. There was a skew towards mutations arising in coding regions with 66% (16/24) mutation targeting coding regions, and 75% (18/24) falling within genes. This is in comparison to 56.8% of the core genome analyzed in this study falling into coding regions, and 63.5% in genes. Assessing the significance of this is complicated by the lower detectability of mutations in non-coding regions that are highly repetitive and have elevated AT content. Restricting analysis to coding SNPs we found 7/9 SNPs resulted in a non-synonymous change, one introduced a premature stop codon and one was a synonymous change. For indels 85.7% (6/7) of coding indels were in frame, and only 1 (14.3%) resulted in a frameshift (Table 1).

**Table 1.**
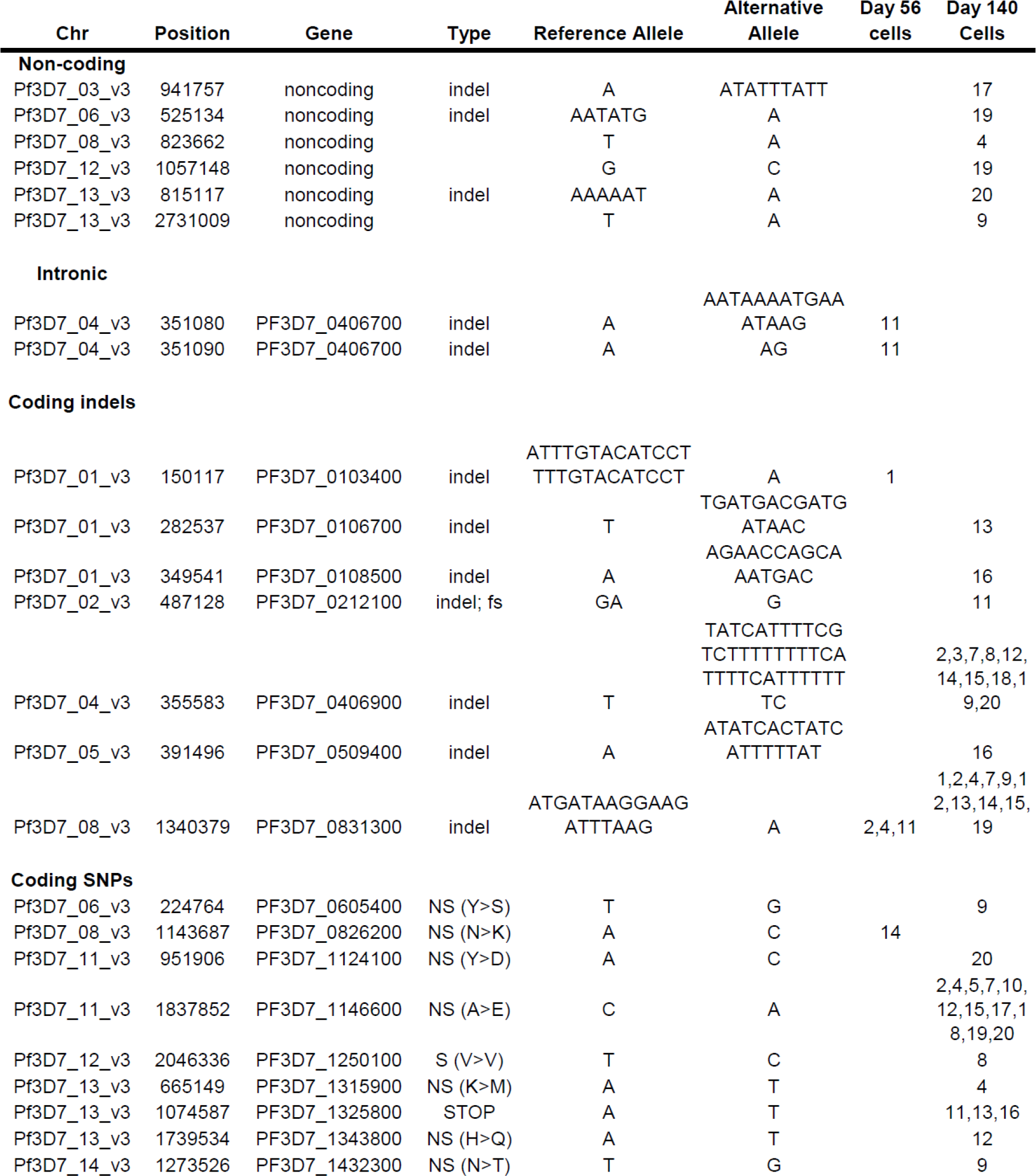
All mutations detected in this study. In the Type column fs denotes a frameshift mutation. The cell ID in which the mutation was discovered is shown in the final columns.

## Discussion

Malaria parasite culture is ubiquitous in malaria research. Our understanding of the basic cell biology of malaria parasites is based heavily on using 3D7, and similarly long term adapted parasite lines. Despite decades in laboratory culture (3D7 was isolated in 1980 (Ponnudurai et al., 1981)) we demonstrate here that adaptive mutations can emerge and rapidly spread through a culture.

### Are these the adaptive mutations?

Both of the mutations identified at high frequency are strong candidates for being a functional mutation. Both occur within genes, and introduce major changes. We cannot discount other mutations that we did not assay with high confidence being the actual drivers. For instance, we did not pursue amplifications or deletions from single cell data. We did not see any evidence of high frequency amplifications or deletions in the bulk sequence. Alternative single cell sequencing protocols are showing promise in identifying copy number variations. We found 15 indels mutated in the dataset. None correlated with the expanded clones. Alternatively, heritable epigenetic changes may have driven the expansion of a clonal lineage, and the mutations we found may be passengers. While feasible, this scenario appears less likely.

### What are adaptive mutations doing?

As we did not pursue the mechanism driving adaptation for these mutations we can only speculate about function. Previous large scale gene knock outs (Bushell et al., 2017, Zhang et al., 2018) have excised PF3D7_1146600 (oocyst rupture protein 1; ORP1) in *P. falciparum* and *P. berghei* and PF3D7_1325800 in *P. falciparum*. For asexual stages, no phenotype was seen and the genes were dispensable. PF3D7_1325800 is annotated as a conserved protein, with no known domains or putative function. Some investigation of PF3D7_1146600 has been performed by others (Curra et al., 2016, Lima et al., 2013). The mutation identified here introduces an amino acid change in the histone-fold domain. This is a critical domain for this protein, it is involved in dimerization of the protein with ORP1. Mutation of the domain prevents oocysts rupturing preventing completion of the parasite life cycle (Curra et al., 2016). This gene showed homology to a group of CCAAT-box binding proteins annotated as NF-YB transcription factors (Lima et al., 2013). However, its sub-cellular location in mosquito stage oocysts is to the oocyst capsule where it appears to play a critical role, rather than in binding DNA. However, PF3D7_1146600 is expressed during the RBC cell cycle suggesting an additional role. Further functional analysis of these genes will be required to define why mutations are adaptive in this setting.

### *In vitro* dynamics support the potential for intrahost competition and local adaptation

Malaria infection with *P. falciparum* can persist for hundreds of days with population sizes in the order of 10^9^−10^12^ parasites. *De novo* mutation will supply millions of potentially beneficial mutations daily (McDew-White et al., 2019). The vast majority of these will be lost by drift before they can be driven to high frequency by positive selection. The dynamics we identify here support the potential for such a process to arise within infections. In addition, the process of adaptation may be accelerated, and its robustness increased by specific population structures which subdivide the population (Pavlogiannis et al., 2018). Such compartmentalization promotes the acquisition of multi-drug resistance in HIV (Moreno-Gamez et al., 2015). Single cell sequencing may allow us to identify mutations segregating at otherwise undetectable frequencies to test these hypotheses.

## Methods

### Cloning and culture of 3D7

3D7 parasites were grown using standard conditions in complete medium (CM) RPMI 1640 with 25mM HEPES, 2.5μg/mL, 50ug/mL hypoxanthine, and 0.5% Albumax II, all from Invitrogen. Cultures were maintained by measuring parasitaemia 3 times a week and carrying them forward at 0.5−2.0% parasitemia in 6mL volume with 4% hematocrit. Cultures were fed 6 days a week, and samples taken biweekly for RBC backups and DNA assays including single cell sorting.

### Flow Cytometry

Cells from culture were pelleted and parasitaemia was measured by thin smear. Seven microliters of packed cells were transferred to 4mL of incomplete media (ICM, RPMI 1640 with 25mM HEPES and 2.5μg/mL all from Invitrogen) pre-warmed to 37°C in a 5mL snap cap polypropylene tube compatible with flow cytometers. Vybrant™ DyeCycle™ Green (Invitrogen) was added to a final concentration of 2uM, cells were mixed by inversion, and placed on a shaker in a 37°C incubator for 30 minutes. After the incubation was completed, the cells were pelleted, supernatant discarded, and washed twice with pre-warmed ICM. Cells were brought up in 2 aliquots of 3mL each with pre-warmed ICM and transferred to the flow cytometer with heat packs.

Cells were sorted with an Influx Cytometer (Becton Dickinson) fitted with a 100μm nozzle using the 488nm laser. Late schizonts were sorted with a 1.0 drop pure setting into 96 well low profile low bind plates (Eppendorf), containing 4uL 1X PBS (Lonza). The plates were centrifuged and immediately placed on dry ice. Plates were then transferred to a −80°C freezer until library processing.

### Single cell sequencing

For library processing, we used the FX Single Cell DNA kit (Qiagen) according to manufacturer directions. Briefly, cells were denatured with D2 buffer at 65°C and the reaction stopped with Stop Buffer. REPLI-g sc DNA polymerase reaction mix was added to the wells at the plate was incubated at 30°C for 2 hours, then stopped by incubation at 65°C for 2 minutes. DNA was measured by QBit, and if concentration was 20ng/uL, the reaction was cleaned and concentrated using a DNA Clean and Concentrate 10 kit (Zymo). The amplified gDNA was fragmented for 33 minutes, then ligated with the adapters included in the Qiagen kit. After ligation the samples were size selected and cleaned using KAPAPure Beads (KAPA). The GeneRead DNA I Amp Kit (Qiagen) was used to amplify 5μL of the libraries to allow for accurate quantitation and size determination. QBit measured DNA content and libraries were pooled to normalize input by concentration. The pooled libraries were measured by the Illumina Library Quant Kit (KAPA) according to manufacturer’s directions, and pool was provided to UT Health Science Center for sequencing on and Illumina NextSeq with a 2×150 flow cell.

### Detection of mutations

We aligned raw sequencing reads to v3 of the 3D7 genome reference (http://www.plasmodb.org) using BWA MEM v0.7.5a (Li, 2013), removed PCR duplicates and reads mapping to the ends of chromosomes (Picard v1.56) and recalibrated base quality scores, realigned around indels and called genotypes using GATK v3.5 (DePristo et al., 2011) in the GenotypeGVCFs mode using QualByDepth, FisherStrand, StrandOddsRatio VariantType, GC Content and max_alterate_alleles set to 6. We recalibrated quality scores and calculated VQSLOD scores using SNP calls conforming to Mendelian inheritance (REF), excluding sites where the VQSLOD score was <0.

A comprehensive description of the protocols in place to prevent contamination and ensure high quality data have been previously published (Nair et al., 2014, Nkhoma et al., 2020, Trevino et al., 2017).

### Phylogenetic reconstruction

We performed Bayesian phylogenetic analysis in BEAST v2.4.3 (Rambaut et al., 2018) with the following parameters: Tip Dates were specified as months with the bulk day 0 was set to 0, day 56 set to 2.0 and day 140 set to 5.0, the GTR substitution model was used with rates estimated from the data, a coalescent constant population was used as the tree prior. We ran the MCMC for 100,000,000 generations, sampling every 2,000. As only variable sites were input into BEAST we edited the ‘constantSiteWeights’ line in the xml file to include the number of non-variable A,C,T and G positions (Kuhner et al., 2000). Convergence was assessed in Tracer v1.6 (Rambaut et al., 2018) and tree were inferred using maximum clade credibility.

### Demographic analysis

Changes in effective population size over time were inferred from a Bayesian skyline plot. We used the same parameters as for construction of a phylogenetic tree, though set the tree prior to ‘Coalescent Bayesian Skyline’. Analysis was implemented in BEAST v2.4.3, and the final skyline reconstruction generated in Tracer v1.6 and plotted in R v3.5.3 (Team, 2017).

## Acknowledgements

We thank Timothy J.C. Anderson for valuable comments and the gift of the 3D7 line used in this study. This study was supported by a NIH grant (NIAID AI110941-01A1 to IHC). I.H.C. is a Milton S. and Geraldine M. Goldstein Young Scientist.

